# Hippocampal subfield volumes contribute to working memory interference control in aging: Evidence from longitudinal associations over 5 years

**DOI:** 10.1101/2023.01.21.525011

**Authors:** P Andersson, G Samrani, M Andersson, J Persson

**Author notes:** Corresponding author: Pernilla Andersson, Center for Life-span Developmental Research (LEADER), School of Behavioral, Social and Legal sciences, Örebro University Fakultetsgatan 1, 702 81, Örebro, Sweden.

## Abstract

In memory, familiar but no longer relevant information may disrupt encoding and retrieval of to-be-learned information. While it has been demonstrated that the ability to resolve proactive interference (PI) in working memory (WM) is reduced in aging, the neuroanatomical components of this decline have yet to be determined. Hippocampal (HC) involvement in age-related decline in control of PI is currently not known. In particular, the association between HC subfield volumes and control of PI in WM has not been examined previously. Here we investigate the associations between mean level and 5-year trajectories of gray matter subfield volumes and PI in WM across the adult life span (N = 157). Longitudinal analyses over 5- years across all participants revealed that reduced volume in the subiculum was related to impaired control of PI. Age-stratified analyses showed that this association was most pronounced in older adults. Furthermore, we found that in older adults the effect of age on PI was mediated by GM volume in the HC. The current results show that HC volume is associated with the ability to control PI in WM, and that these associations are modulated by age.

## 1. INTRODUCTION

The constant accumulation of memory representations places high demands on a cognitive system that is efficient in controlling the content of memory in order to prevent interference from goal-irrelevant information. Proactive interference (PI) arises from existing memory representations conflicting with new target information. While PI has most commonly been investigated within the domain of episodic memory (Keppel, 1968; Roediger & McDermott, 1995), there is now much evidence that PI plays a critical role also in working memory (WM; Bunting, 2006; Emery, Hale, & Myerson, 2008). PI is also closely linked with performance levels on WM span tests (Kane & Engle, 2000; Lilienthal, 2017; Lustig, May, & Hasher, 2001; May, Hasher, & Kane, 1999).

While there is much evidence that WM declines with increasing age, evidence for age-related impairments in inhibition and interference control has been less consistent (Rey-Mermet & Gade, 2017; Verhaeghen, 2011). In general, it seems that in tasks in which conflicting information can be handled within the focus of attention show less of a decline compared to tasks in which the conflict arises from familiar, but no longer relevant information outside the focus of attention (Lustig & Jantz, 2014). One possibility is that age-related cognitive decline is mediated by deficits in interference control or other executive functions, with corresponding consequences on other aspects of cognition (Hasher & Zacks, 1988; McGabe, Roediger, McDaniel, Balota, & Hambrick, 2010). Indeed, an age-related decline in PI was recently demonstrated using a population-based life-span sample of adults (Samrani & Persson, 2021), and in this particular study PI was also found to predict mean level performance in other cognitive abilities.

Age-related impairments in interference control are most likely related to less efficient brain function, and changes in brain morphology and neuromodulation that occur with increasing age. Much of the work on the neural bases of interference control in WM has focused on the prefrontal cortex. There is a general consensus that the left inferior frontal gyrus, together with the anterior cingulate cortex, and the insula are key regions involved in successful resolution of PI in WM (Badre & Wagner, 2005; Burgess & Braver, 2010; Henson, Shallice, Josephs, & Dolan, 2002; Jonides & Nee, 2006; Marklund & Persson, 2012; Nelson, Reuter-Lorenz, Persson, Sylvester, & Jonides, 2009; Persson, Larsson, & Reuter-Lorenz, 2013; Samrani, Bäckman, & Persson, 2019). However, despite evidence suggesting that the hippocampus (HC) is critical for interference control (Öztekin, Curtis, & McElree, 2008), little is known on how these regions contribute to resolving PI in WM. This is particularly important in studies on aging given the well documented effects of aging on HC gray matter volume (Gorbach et al., 2017; Persson et al., 2012; Raz, Rodrigue, Head, Kennedy, & Acker, 2004; Raz et al., 2003).

While the HC has been the focus of research of long-term memory functions for decades, its role in WM is less understood. There are indications that the HC may contribute to WM processes; activation in the HC has been demonstrated during fMRI WM tasks (Axmacher, Schmitz, Weinreic, Elger, & Fell, 2008; D E Nee, Jonides, & Berman, 2007), and patients with HC lesions are impaired in WM tasks (Cashdollar et al., 2009; Hannula, Tranel, & Cohen, 2006; Nichols, Kao, Verfaellie, & Gabrieli, 2006). There is also some recent functional imaging evidence showing that the HC is involved in successful resolution of PI (Öztekin et al., 2008). In particular, the HC may play a key role in distinguishing older from newer information in long term memory (Barredo, Öztekin, & Badre, 2015; Caplan, McIntosh, & De Rosa, 2007; Yassa & Stark, 2011), and WM (Leszczynski, 2011; Nauer, Whiteman, Dunne, Stern, & Schon, 2015; Ranganath & D’Esposito, 2001; Öztekin, Davachi, & McElree, 2010) thereby reducing PI. The HC could also be involved in resolving PI by contributing context or source information that can help distinguish familiar but no longer goal-relevant items from goal-relevant target items. This key function may be critical for both long-term memory and WM.

In contrast to episodic memory, the neuroanatomical underpinnings of interference control have received much less attention despite its likely role in age-related cognitive impairments. To the best of our knowledge, only one study to date has examined the cerebral morphological properties underlying interference control across the adult life span using both cross-sectional and longitudinal data (Samrani et al., 2019), and this particular study focused on frontal regions only. We focus here on the HC because of their known role in memory functions. In the current study, we use data from a population-based cohort study to examine cross-sectional (N = 280) and longitudinal (N = 157) associations between HC subfield gray matter, and PI in WM across the adult life span. We hypothesize that larger gray matter volume in the HC should be related to better interference control (i.e., less PI). Since it has been demonstrated that brain function and volume might be differentially linked to cognition in younger and older adults (Koen & Rugg, 2019; Park et al., 2004; Rieckmann, Johnson, Sperling, Buckner, & Hedden, 2018; Van Petten, 2004), and that brain volume – cognition interactions may be stronger in older adults (Burzynska et al., 2012) we performed age-stratified analyses within groups on younger/middle-aged and older adults, as well as across the whole sample. We also obtained HC subfield volumes to investigate potential differential involvement of different sub-fields. We believe that this study takes critical steps to advance our understanding of hippocampal contributions to interference control across the adult life span.

## 2. MATERIAL AND METHODS

### 2.1 Participants

Participants were recruited from *The Betula prospective cohort study: Memory, health, and aging* (Nilsson et al., 1997), a deeply phenotyped longitudinal cohort. Participants were included from samples for which MRI measures were collected in 2008-2010 and 2013-2014. The age of the participants at baseline ranged from 25 to 90 years (mean = 56.9, standard deviation [SD] = 12.1; 49% female), and mean education level was 13 years (SD = 4.0 years). Individuals with clinical dementia and other neurological disorders at baseline were excluded as were participants with missing longitudinal data. Participants were screened with the Mini-Mental State Examination (MMSE; Folstein, Folstein, & McHugh, 1975), and those scoring 25 and above at both baseline and follow-up were included. Participants with extremely low performance on the n-back task (proportion hits minus proportion false alarms < .1), indicating a very low adherence to task instructions, were not included. Thus, the total sample consisted of 157 participants. For a drop-out analysis, see Nyberg et al., (2019). To retain the diversity of the sample, exclusions were not made for: diabetes, hypertension, mild depressive symptoms, and other moderately severe medical conditions, which are common among the elderly. The Betula study was approved by the Regional Ethical Review Board in Umeå, and written consent was obtained from every participant. We investigated whether age modulated the relationship between brain volume and PI by dividing participants into two groups: one consisting of younger and middle-aged adults (25-65 years at baseline), and one consisting of older adults (70-90 years at baseline). A complementary analysis was also performed by dividing participants into three groups: one consisting of younger adults (25-45 years), one consisting of middle-aged adults (50-60 years) and one consisting of older adults (65-85). The number of participants within different age groups can be found in Supplementary Figure 1. Demographic information and cognitive performance can be found in Table 1.

**Table 1.**
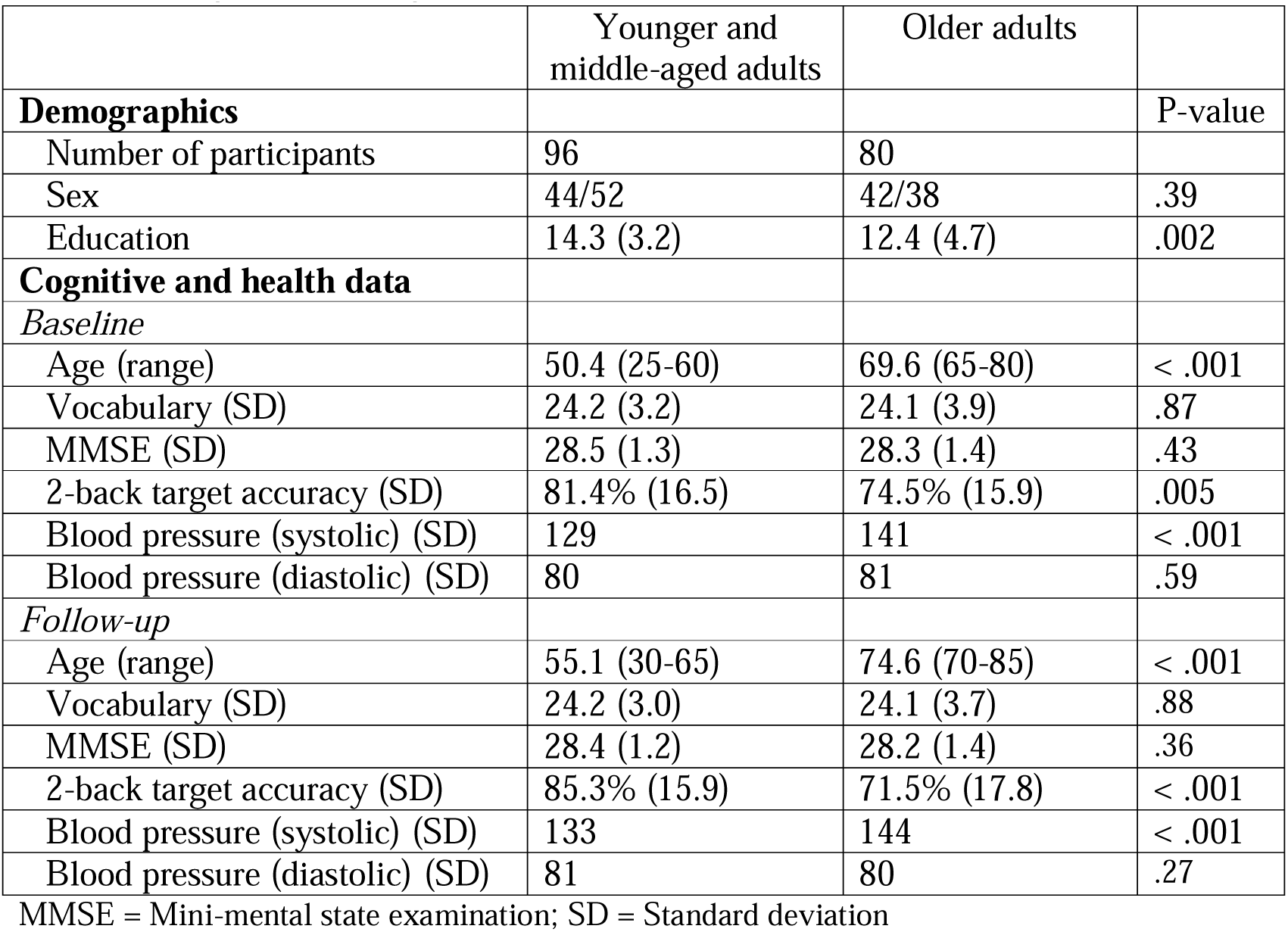
Demographic and cognitive measurements

|  | Younger and middle-aged adults | Older adults |  |
| --- | --- | --- | --- |
| **Demographics** |  |  | **P-value** |
| Number of participants | 96 | 80 |  |
| Sex | 44/52 | 42/38 | .39 |
| Education | 14.3 (3.2) | 12.4 (4.7) | .002 |
| **Cognitive and health data** |  |  |  |
| *Baseline* |  |  |  |
| Age (range) | 50.4 (25-60) | 69.6 (65-80) | < .001 |
| Vocabulary (SD) | 24.2 (3.2) | 24.1 (3.9) | .87 |
| MMSE (SD) | 28.5 (1.3) | 28.3 (1.4) | .43 |
| 2-back target accuracy (SD) | 81.4% (16.5) | 74.5% (15.9) | .005 |
| Blood pressure (systolic) (SD) | 129 | 141 | < .001 |
| Blood pressure (diastolic) (SD) | 80 | 81 | .59 |
| *Follow-up* |  |  |  |
| Age (range) | 55.1 (30-65) | 74.6 (70-85) | < .001 |
| Vocabulary (SD) | 24.2 (3.0) | 24.1 (3.7) | .88 |
| MMSE (SD) | 28.4 (1.2) | 28.2 (1.4) | .36 |
| 2-back target accuracy (SD) | 85.3% (15.9) | 71.5% (17.8) | < .001 |
| Blood pressure (systolic) (SD) | 133 | 144 | < .001 |
| Blood pressure (diastolic) (SD) | 81 | 80 | .27 |

MMSE = Mini-mental state examination; SD = Standard deviation

### 2.2 Cognitive measures

PI was measured using a verbal 2-back WM task which included familiar lure items (Figure 1, Gray, Chabris, & Braver, 2003; Marklund & Persson, 2012; D E Nee et al., 2007) occurring either 1 or 2 trial(s) after the target position. The task included 40 trials; 21 non-familiar trials, 9 target trials, 8 3-back lures, and 2 4-back lures. *Target trials* matched the same stimuli as presented two trials earlier and required a ‘Yes’ response. *Lure trials* consisted of stimuli already presented 3- or 4 trials earlier and required a ‘No’ response. *New trials* were non-familiar trials that had never been presented and required a ‘No’ response. Stimuli and trial conditions were presented in the same fixed order for all participants. Stimuli consisted of Swedish nouns and were presented one at a time for 2.5 s, with an inter trial interval of 2 s. For each presented word, participants were instructed to press the “m” key on a standard Swedish keyboard, which corresponds to ‘Yes’ (“Yes, the word I now see has been shown two words ago”) and the “x” key for ‘No’ (“No, the word I now see has not been shown two words ago”). Participants were instructed to answer as quickly and accurately as possible.

**Figure 1.**
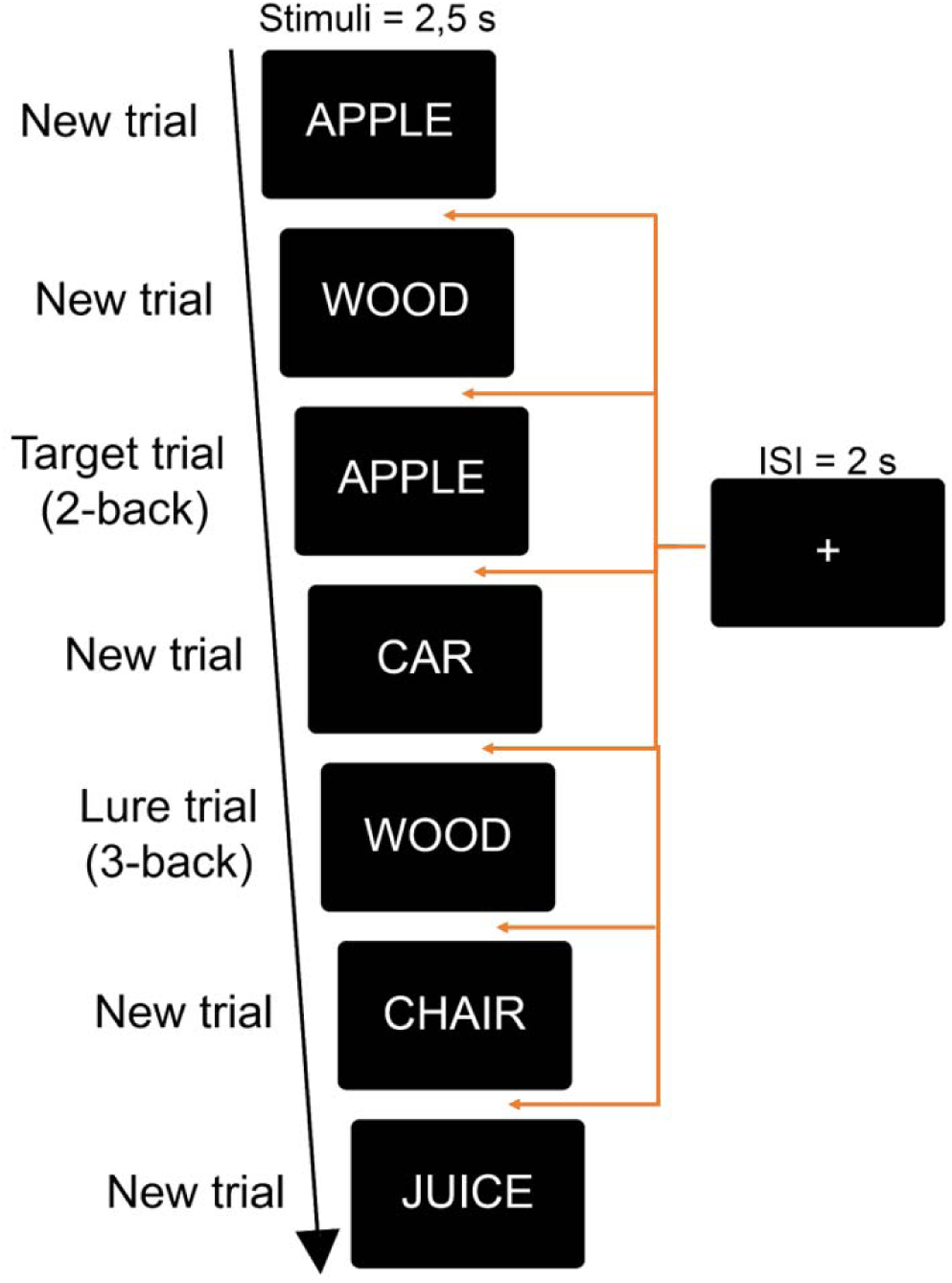
N-back task. Experimental design and trial organization

PI scores were calculated by combining the relative proportional difference in RT and accuracy between non-familiar trials (new trials) and 3- and 4-back familiar trials (lure trials). The PI scores based on accuracy and RTs were positively correlated, both at baseline (r = .295, P < 0.0005) and follow-up (r = .372. P < 0.0005), indicating shared variance between the two measures of PI. Median RTs were used to reduce the influence of extreme values. Accuracy data for working memory performance in the n-back task is also reported under section 3.1.

### 2.3 MRI data acquisition and analyses

The MRI data was collected using a 3T GE scanner, equipped with a 32-channel head coil. The same scanner was used for baseline and follow-up data collection. T1-weighted images were acquired with a 3D fast spoiled gradient echo sequence (180 slices with a 1 mm thickness; TR: 8.2 ms, TE: 3.2 ms, flip angle: 12 degrees, field of view: 25 x 25 cm).

To extract HC volumes, T1-weighted images were processed using the Freesurfer image analysis suite. Hippocampal subfield segmentation was performed using the longitudinal stream (Reuter et al., 2012) of the Hippocampal Subfield module of FreeSurfer v.6.0 (https://surfer.nmr.mgh.harvard.edu/fswiki/HippocampalSubfields). The technical details of these procedures are described in prior publications (Dale, Fischl, & Sereno, 1999; Fischl et al., 2002; Fischl et al., 2004; Jovicich et al., 2006). Automated cortical and subcortical parcellation tools in the FreeSurfer software were used for volumetric segmentation, cortical surface reconstruction, and parcellation to quantify the brain volumes of interest. Cortical reconstructions and volumetric segmentations were performed on all images by executing a semi-automatic processing step (recon-all) within this software (Dale et al., 1999; Fischl et al., 2002). Identification of hippocampal subfields using FreeSurfer 6.0 should be considered probabilistic, since it uses prior knowledge from ex vivo brains scanned at 7 T MRI combined with the available contrast information from MR images (Iglesias et al., 2016).

### 2.4 Scanner stability

From baseline to follow-up data collection, changes were implemented to the scanner software, and it is critical to check that these did not affect image quality. A quality assurance program based on Friedman and Glover (2006) was therefore run on a weekly basis on the scanner. For the structural data, the same T1-weighted fast spoiled gradient echo protocol as in the study was used to obtain volume data for the GE phantom. A threshold well above the noise level and the selected voxels were used to calculate the volume of the phantom. The relative volume change between the beginning of the quality assurance measurements and the time for T6 data collection was 0.45%, which is small compared to the expected average volume change in cortical regions in healthy elders (Fjell et al., 2009). Furthermore, the positive change in the measured volume of the phantom suggests that any significant volumetric decline observed in the study might be slightly underestimated.

### 2.5 Definition of cognitive and volumetric change

Change in proactive interference was estimated by dividing the PI-score from follow-up by the baseline PI-score, with positive values indicating increase in PI and negative values indicating decrease in PI over time. Similarly, we measured change in brain volumes by dividing the values from the second time point by the values from the first time point. The relative change from time points 1 to 2 takes into account the brain size for each individual, thus controlling for brain size without including TIV as a covariate in the analysis. Significant negative correlations between baseline level and rate of change in proactive interference were observed in both age cohorts (Pearson’s *r* were -0.574 for younger/middle-aged adults and - 0.704 for older adults). Individuals with low initial performance tended to improve or decline less rapidly and individuals starting with high levels of performance seemed to display more decline.

### 2.6 Selection of regions of interest

In order to reduce the number of comparisons, and because we did not have any specific hypotheses regarding laterality, we averaged volumes of the left and right hemisphere. For HC subfield analyses, we included the combined left and right CA1, dentate gyrus (DG), CA3, and subiculum subfields. We also performed analyses using total HC volume.

### 2.7 Statistical analyses

In the current study, relationships between cross-sectional estimates of brain volume and PI, as well as brain markers of change and change in PI were assessed using ordinary and partial correlation coefficients. Associations were controlled for age and education. Cross-sectional volumetric analyses were controlled for total intracranial volume (TIV). Cross-sectional analyses were performed on the average PI and HC volume of baseline and follow-up. Longitudinal analyses were additionally controlled for baseline level of PI. Partial eta squared (□_p_^2^) was used to measure effect size. Statistical analyses were performed using SPSS software ver. 26.0 (IBM, Armonk, NY, USA). Analyses were first performed across the whole age span, but since the relationship between local brain volume and cognition might change with increasing age (Burzynska et al., 2012; Razlighi et al., 2017), we also tested for associations within each of the two age groups. Multivariate outliers were identified using Mahalanobis distance at a P < .001.

In addition to analyses across the whole sample, we also tested whether there are age differences in the GM volume – PI associations. To do this, and similar to other studies (e.g. Andersson, Li, & Persson, 2022; Razlighi et al., 2017), we stratified participants into one group of younger and middle-aged (age at baseline: 25-65), and one group of older (age at baseline: 70-85) adults. However, we also performed age-stratified analyses using one group of younger adults (age at baseline: 25-45), one group of middle-aged adults (age at baseline: 50-60), and one group of older adults (age at baseline: 65-85).

Moderation and mediation analyses were performed using the PROCESS macro (Hayes, 2013) implemented in SPSS. Using moderation analyses, we examined whether age was a significant moderator of the relationship between HC volume and PI. Mediation analyses were carried out to test whether the relation between the predictor (age) and the outcome (PI) was mediated – in total or in part – by a mediator variable (HC volume). For both moderation and mediation analyses, a bootstrapping resampling strategy was implemented using 5000 bootstrap samples. For the mediation analysis, path *a* describes the direct effect of age on HC volume, path *b* represents the direct effect of HC volume on PI, and path *c* represents the direct effect between age and PI. Finally, path *c*′ indicates the total effect of age on PI when HC volume is included in the model (Figure 3). Bias-corrected 95% confidence interval (CI) was computed to evaluate the contribution of the mediator (indirect effect, path a × b). CI reached significance when the interval range did not include zero. In order to examine the hypothesis that mediation would only be present in older adults, separate mediation analyses were carried out in younger/middle-aged and older adults, as well as in the whole sample. Standardized coefficients are reported for all mediation analyses.

**Figure 2.**
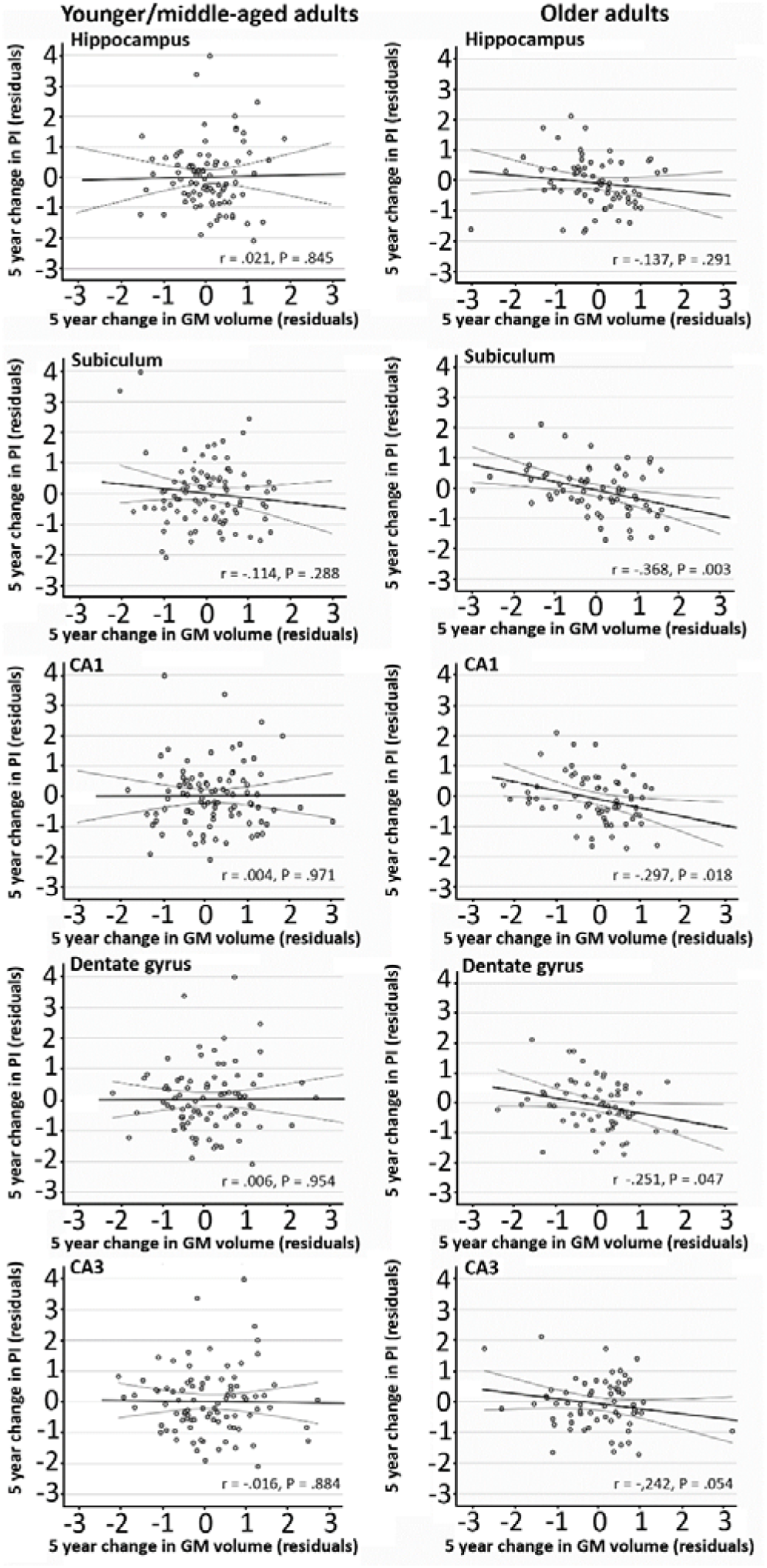
Correlations between relative interference score and GM volume. (Top) Correlations between relative interference score and HC volume at baseline for young/middle- aged and older adults. Standardized residuals are corrected for age, education, and TIV. (Bottom) Correlations between relative interference score and HC sub-field volumes (CA1 and CA3/DG) at baseline for young/middle-aged and older adults. Standardized residuals are corrected for age, education, and TIV.

**Figure 3.**
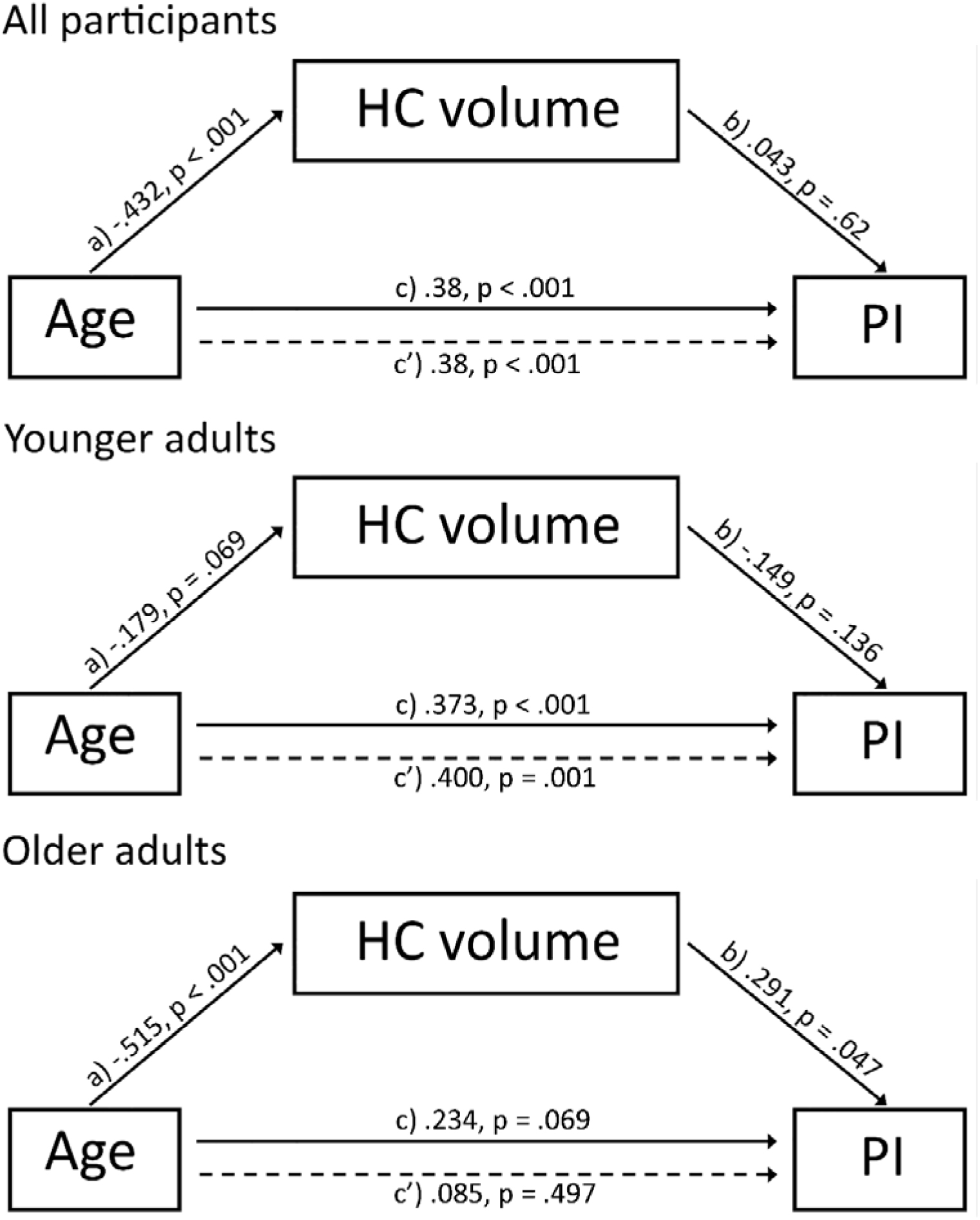
Mediation analysis. Mediation analysis investigating the three-way association between age, HC volume and PI. (A) Mediation analysis (model A) where “a” indicates the effect of age on HC volume, “b” indicates the effect HC volume on PI, “c” indicates the direct effect of age on PI, and “c’” indicates the total effect (direct and indirect) of age on PI.

## 3. RESULTS

### 3.1 Older age was associated with higher levels of proactive interference

Accuracy for non-familiar lures (98%) was significantly higher than for both familiar lures (63%; F(175) = 110, p < .001, □_p_^2^ = .381) and targets (78%; F(175) = 264, p < .001; □_p_^2^ = .597) and is on par with other studies using the modified version of the n-back task to induce familiarity based PI (Loosli, Rahm, Unterrainer, Weiller, & Kaller, 2014; Samrani, Bäckman, & Persson, 2017; Samrani & Persson, 2021) in older adults. Similarly, RT for non-familiar lures (1011 ms) was faster than for both familiar lures (1424ms; F(175) = 311, p < .001, □_p_^2^ = .639) and targets (1101 ms; F(175) = 19.4, p < .001, □_p_^2^ = .098).

Moreover, older individuals had a lower ability to control interference as indicated by the positive correlation between age and PI (r = .365, P < .0001). .The relationship between change in PI over 5 years and age was, however, not significant (P > .1).

### 3.2 Cross-sectional and longitudinal effects of age on hippocampal brain volume

Linear regression analyses showed that higher age was a significant predictor of lower total HC volume (r^2^ = .323, β = -.437, P < .0001). Similarly, analyses of HC sub-field volumes showed that age was a significant predictor of CA1 volume (r^2^ = .253, β = -.255, P < .001) CA3 volume (r^2^ = .331, β = -.189, P = .007), DG volume (r^2^ = .34, β = -.303, P < .001) and subiculum volume (r^2^ = .262, β = -.377, P < .001).

### 3.3. Cross-sectional associations between hippocampus volume and proactive interference in working memory

For the analyses across the whole sample, the relationship between mean level PI and total HC volume was not significant (r =.038, *p* =.62). Moreover, none of the associations with HC subfield volume were significant (all Ps > .1).

When analyses were performed for each age group separately, we found that in younger and middle-aged adults, none of the associations between PI and HC volume were significant (all Ps > .1). In the group of older adults, with the association between total HC volume and PI approached significance (r = .232*, p* = .05), but none of the associations between HC subfields and PI were significant (all Ps > .1).

### 3.4. Longitudinal associations between hippocampus volume and proactive interference in working memory

The correlation between 5-year change in HC volume and PI across the whole sample was not significant (r = -.037, P = .65). However, analyses on sub-field volume showed a significant, negative, correlation between change in subiculum volume and change in PI (r = -.213, P = .009) with decreased subiculum volume correlating with increased PI. All other correlations were non-significant (P > .1). Age-stratified analyses for the total HC volume were non-significant for both younger/middle-aged and older adults (P > .1), but analyses on sub-field volumes showed that in the group of older adults, change in HC sub-field volume was negatively associated with change in PI (Figure 2; CA1: r = -.297, P = .018; Subiculum: r = -.368, P = .003; DG: -.251, P = .047) with decreased sub-field volume correlating with increased PI. Moderation analyses showed that age was not a significant moderator for the relationship between change in HC sub-field volume and change in PI. However, for the subiculum, results from the Johnson-Neyman procedure showed that this relationship was significant in participants above 58 years of age.

### 3.5 Complementary analysis with three age groups

Because of the large age range in the younger/middle-aged age group, we also performed analyses in which we divided the sample into three age groups (see Methods). Note that the older age group was similarly defined as in the two-groups analyses. We did not observe any significant differences in HC volume – PI associations between the younger and middle-aged age groups, neither in the cross-sectional nor the longitudinal analyses. This further indicates that the observed associations between 5-year change in HC volume and PI in working memory were restricted to the group of older adults.

### 3.6 Mediation analyses

Single mediation analysis across all participants showed that HC volume was not a significant mediator of the effect of age on PI (estimated effect: -.018; lower level (LL) CI: -.118, higher level (HL) CI: .062). Direct effects between age and HC volume (b = -.432, LLCI: -23.0, HLCI: -12.3P < .001) and age and PI (b = .38, LLCI .012, HLCI: .029, P < .001) were significant, but the effect between HC volume and PI was not significant (b = .043, LLCI: -.001, HLCI: .001, P = .62). Age-stratified mediation analysis showed that for younger and middle-aged adults HC volume was not a significant mediator of the effect of age on PI (estimated effect: .027; LLCI: -.018, HLCI: .089). The direct effect between age and PI was significant (b = .373, LLCI: .012, HLCI: .038, P < .001), but the direct effects between and age and HC volume (b = -.179, LLCI -15.6, HLCI: .619, P = .069) and HC volume and PI (b = -.149, LLCI: -.001, HLCI: .001, P = .136) were non-significant. For older adults, however, HC volume was a significant mediator of the effect of age on PI (estimated effect: -.15; LLCI: -.304, HLCI: -.021). The direct effects between age and HC volume (b = -.515, LLCI: -79.1, HLCI: -36.9, P < .001) and HC volume and PI (b = .291, LLCI: .001, HLCI: .007, P = .047) were significant, but not the direct effect between age and PI (b = .234, LLCI: -.006, HLCI: .069, P = .094)

## 4. DISCUSSION

There is much evidence from patient and functional neuroimaging studies implicating HC as critical for efficient resolution of PI in WM. The current findings confirm the importance of the HC in interference control and suggest that this region is differentially involved in PI resolution in younger and older adults. Cross-sectional analyses did not reveal any significant association between HC volume or sub-field volume in the whole sample or in age-stratified groups. Across all participants, HC volume was not significantly associated with PI. Longitudinal analyses showed that in older adults, 5-year decrease in subiculum, CA1, and DG volume was related to decreased control of PI over 5 years. Finally, age-stratified mediation analyses revealed that, in older adults, whole HC volume significantly mediated the age – PI relationship. These results provide important insights into the neural architecture underlying interference control in WM.

Based on previous findings of a negative association between frontal cortex volume and PI (Samrani et al., 2019), along with data from fMRI of a link between HC activation and PI (D E Nee et al., 2007; Samrani & Persson, 2022; Öztekin et al., 2008), we expected that larger HC volume would be linked to less PI. Cross-sectional analyses of the relationship between HC volume and PI did not reveal any significant associations for the whole sample or for the separate age-groups. Likewise, sub-field volume was not found to be significantly associated with PI.

Longitudinal analyses of change across 5 years demonstrated that decreased HC volume was associated with decreased control of PI across all participants, but only in the subiculum. Age-stratified analyses found that decreased volume in the CA1, subiculum and DG subfields, respectively, was significantly correlated with concurrent increase in PI in the older group. These findings corroborate previous cross-sectional findings that demonstrate a relationship between hippocampal function and PI in long-term memory (Blumenfeld & Ranganath, 2007; Wanjia, Favila, Kim, Molitor, & Kuhl, 2021). Moreover, it was recently demonstrated that 4 weeks of mindfulness training both improved the ability to control interference in WM and also resulted in an increase in HC volume (Greenberg et al., 2018). Importantly, the authors also found that mindfulness-related increase in HC volume was related to reduced PI.

In the older group, decreased sub-field volume in the CA1, DG, and subiculum, respectively, was related to increased PI over five years. The contribution of different HC sub-fields to memory functioning is not fully understood yet and most research has focused on the contributions to long-term memory processes.

Specific subfields within the HC have been linked to pattern separation and pattern completion processes. While the n-back task used in the present study was not designed to measure pattern separation, the similar context across stimuli may require the engagement of such, or similar, processes in order to overcome PI during lure trials. The DG and CA3 sub-fields being differentially involved in pattern separation processes (Azab, Stark, & Stark, 2014; Berron et al., 2016; Stevenson et al., 2016; Yassa, Stark, et al., 2011) and the CA1 subfield has been linked to pattern completion processes (Bakker et al., 2008). However, some divergent results have also been reported such as CA3 being implicated in pattern completion in episodic memory (Guzowski, Knierim, & Moser, 2004; Rolls, 2016). The involvement on the subiculum in these processes has not been discerned as a relationship has been observed with both pattern separation (Bouyeure et al., 2021; Potvin et al., 2009) and completion (Bakker et al., 2008). In a recent study, focusing on sub-field contribution to separation/completion processes in working memory, increased activity in the CA3/DG, subiculum, and CA1 was observed in relation to pattern separation while pattern completion was associated with increased activity in the CA1, subiculum, and entorhinal cortex (Newmark et al., 2013). Thus, complementing findings from studies on long-term memory. Clearly, more specific investigations using paradigms targeting pattern separation and completion processes would need to be conducted to draw any conclusions on possible relationship with processes involved in interference control.

While the subiculum has received less attention than hippocampal subfields commonly considered part of the hippocampus proper, recent studies suggest that this region may play an important role in memory (e.g., Hartopp et al., 2019; Ku et al., 2017). For example, the region has been implicated in recognition memory and task memory load (Ku et al., 2017).

Interestingly, in a recent study, Hartopp and colleagues (2019) found significant associations between subiculum volume and fornix microstructure in a group of 31 healthy, older adults. Furthermore, this association was found to significantly contribute to episodic memory performance, suggesting that age-related alterations in subiculum structure could be driving fornix microstructural degradation. In another study, a similar observation was made of the DG mediating the effect of reduced fornix integrity on the retrieval of learned associations (Hayek et al., 2020). Similar investigations of the relationship between changes in fornix microstructure and volumetric changes in HC sub-fields have, to our knowledge, not been conducted in relation to working memory. However, a recent study investigating correlations between white matter integrity and control of PI in working memory found that reduced fornix FA was associated with reduced ability to control PI across five years (Andersson et al., 2022). Follow-up analyses combining volumetric and microstructural measures could shed light on whether fornix – HC sub-field associations also contribute to the ability to control PI in working memory.

It is important to emphasize that we do not imply that the HC is uniquely involved in interference control. Rather, we believe that this region belongs to a network of regions, including the inferior frontal gyrus, anterior cingulate cortex, and insula that are all involved in control of PI. For example, animal tract-tracing studies have demonstrated direct connection between the HC and prefrontal cortex (Pandya, Van Hoesen, & Mesulam, 1981; Suzuki & Amaral, 1994) and anterior cingulate cortex (Morris, Pandya, & Petrides). Moreover, numerous human diffusion-tensor imaging (Draganski et al., 2008) and functional connectivity (D E Nee & Jonides, 2014) studies provide evidence for HC connectivity with the prefrontal cortex.

The current study has some notable limitations that need to be mentioned. First, our longitudinal analyses contained only data from two measurement points. Therefore, we were not able to examine long-term trajectories of change, and this design also did not permit independent estimations of retest effects, which are known to influence longitudinal data. Future studies that include three or more time points could provide data on long-term trajectories of change that we are unable to show in the current study design and may also provide more accurate estimation of retest effects. Second, the relatively low number of participants that remained for follow-up testing may have resulted in low statistical power in detecting longitudinal effects. This also restricted our age-differential analyses to include only cross-sectional data since the within-group sample size was too small for longitudinal analyses.

Taken together, our results support the idea that the hippocampus plays a noticeable role in controlling PI in WM. Additionally, we demonstrate that the HC is differentially involved in PI resolution in younger and older adults. We provide new evidence that reduced HC subfield volumes over 5-years was related to concurrent increase in PI in older adults. Furthermore, we demonstrate that HC volume significantly mediates the relationship between age and PI in older, but not younger/middle aged adults. We believe that these results provide new and important insights into the neural architecture underlying interference control in WM.

## 5. AUTHOR CONTRIBUTION

P.A: Writing - Original Draft Preparation, Writing - Review & Editing

G.S: Investigation, Writing - Original Draft Preparation

M.A: Formal Analysis, Investigation,

J.P: Conceptualization, Formal Analysis, Writing - Original Draft Preparation, Writing - Review & Editing

## 6. SUPPLEMENTARY MATERIAL

Supplementary figure 1. Distribution of age across sample

## 7. ACKNOWLEDGEMENTS

We acknowledge the contribution by the staff in the Betula project and all participants. The Freesurfer analyses were performed on resources provided by the Swedish National Infrastructure for Computing (SNIC) at HPC2N in Umeå.

## 10. DATA AVAILABILITY

The data that support the findings of this study are available from the corresponding author upon reasonable request.

