## Supplementary figures and images for "Hippocampal subfield volumes contribute to working memory interference control in aging: Evidence from longitudinal associations over 5 years"

### Supplementary Figure 1

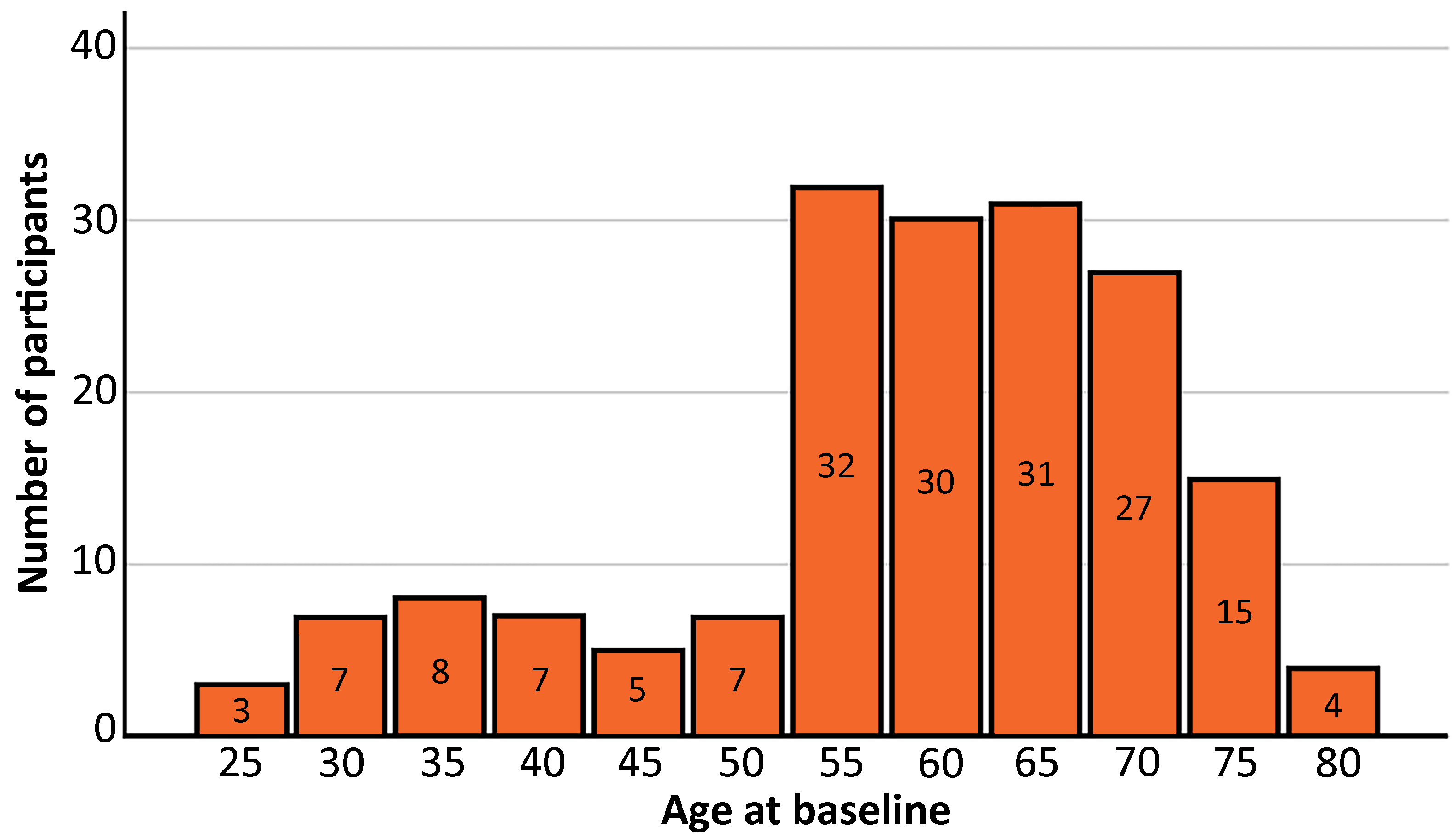
